# M1 disruption delays motor processes but not deliberation about action choices

**DOI:** 10.1101/501205

**Authors:** Gerard Derosiere, David Thura, Paul Cisek, Julie Duque

**Author notes:** Corresponding author contact details: **Gerard Derosiere**, CoActions Lab, Institute of Neuroscience, Université catholique Louvain, Av. Mounier, 53 - Bte B1.53.04, 1200 Bruxelles, Belgium.

## Abstract

Decisions about actions typically involve a period of deliberation that ends with the commitment to a choice and the motor processes overtly expressing that choice. Previous studies have shown that neural activity in sensorimotor areas, including the primary motor cortex (M1), correlates with deliberation features during action selection. Yet, the causal contribution of these areas to the decision process remains unclear. Here, we investigated whether M1 determines choice commitment, or whether it simply reflects decision signals coming from upstream structures and instead mainly contributes to the motor processes that follow commitment. To do so, we tested the impact of a disruption of M1 activity, induced by continuous theta burst stimulation (cTBS), on the behavior of human subjects in *(1)* a simple reaction time (SRT) task allowing us to estimate the duration of the motor processes and *(2)* a modified version of the tokens task (Cisek et al., 2009), which allowed us to estimate subjects’ time of commitment as well as accuracy criterion. The efficiency of cTBS was attested by a reduction in motor evoked potential amplitudes following M1 disruption, as compared to those following a sham stimulation. Furthermore, M1 cTBS lengthened SRTs, indicating that motor processes were perturbed by the intervention. Importantly, all of the behavioral results in the tokens task were similar following M1 disruption and sham stimulation, suggesting that the contribution of M1 to the deliberation process is potentially negligible. Taken together, these findings favor the view that M1 contribution is downstream of the decision process.

**New and noteworthy:** Decisions between actions are ubiquitous in the animal realm. Deliberation during action choices entails changes in the activity of the sensorimotor areas controlling those actions, but the causal role of these areas is still often debated. Using continuous theta burst stimulation, we show that disrupting the primary motor cortex (M1) delays the motor processes that follow instructed commitment but does not alter volitional deliberation, suggesting that M1 contribution may be downstream of the decision process.

## INTRODUCTION

The physical world provides animals with a variety of action opportunities, constantly requiring them to make decisions, some of which are critical for survival. For instance, the choice of a car driver to turn left or right in front of a sudden obstacle may have dramatic consequences on her/his life and on that of the pedestrians around. The driver will have to quickly deliberate and commit to one action.

Deliberation about actions is thought to entail a competition between distinct neural populations within the motor system (Pezzulo and Cisek, 2016; Svoboda and Li, 2018). In this view, separate action opportunities increase activity of distinct populations, which compete against each other, possibly through mutual inhibition (Michelet et al., 2010). An action is eventually selected and executed when activity in the related population reaches a critical decision threshold (Laming, 1968; Ratcliff, 1978; Stone, 1960).

In line with this hypothesis, a compendium of studies has shown that the dorsal premotor (PMd), but also the primary motor cortex (M1), display a buildup of choice-selective activity during the decision process. The rate of this buildup depends on the amount of sensory evidence favoring the selection of each action in the environment (Alamia et al., 2018; Derosiere et al., 2018; Donner et al., 2009; Gould et al., 2012; Tosoni et al., 2014; Wyart et al., 2012). According to this view, in the car driver example above, the presence of pedestrians on the right side of the street would increase the activity of the population coding for the movement of rotating the wheel towards the left, and possibly weaken the activity of the population favoring the opposite rightward rotation movement. Ultimately, the driver will commit to turning left and execute the related action, to avoid hitting the group of people.

Importantly, making decisions often requires balancing the desire to take time to deliberate accurately (*i.e.*, to accumulate sensory evidence and make the best choice) with the urge to act (Churchland et al., 2008; Forstmann et al., 2008; Hanks et al., 2011; Thura and Cisek, 2014a). During a speeded decision, the urge to act increases as time passes (Cisek et al., 2009, Ditterich, 2006, Drugowitsch et al., 2012, Seideman et al., 2018; Thura et al., 2012) but the overall level of urgency also varies depending on the context (Murphy et al., 2016, Thura et al., 2014; Thura and Cisek, 2016, 2017). In the situation described earlier, the driver’s level of urgency will be higher if the obstacle suddenly appears close to the car than if it appears far away. As evident in this example, adjustments in urgency alter the balance between decision speed and accuracy - *i.e.*, the so-called speed-accuracy tradeoff: choices are more likely to be incorrect when time pressure is elevated, while accuracy improves when temporal demands permit long deliberation (*e.g.*, Hanks et al., 2011; Seideman et al., 2018).

At the neural level, several lines of evidence indicate that higher levels of urgency during deliberation about action choices modulate neural activity in PMd and M1 (Murphy et al., 2016; Steinemann et al., 2018, Thura and Cisek, 2014a, 2016). In these areas, activity is globally amplified at baseline and then builds-up at a faster rate when urgency is high compared to when it is low, reducing the time needed to reach decision threshold but at the cost of accuracy (Thura and Cisek, 2016). Recent findings suggest that the basal ganglia (Thura and Cisek, 2017; van Maanen et al., 2016) and the locus coereleus (Hauser et al., 2018; Murphy et al., 2016) may contribute to generate such a modulation of motor cortical activity.

Together, these data indicate that the motor cortical areas combine both the sensory evidence signals guiding the choice with the urgency-related signals determining the best time to commit to that choice, suggesting a crucial role of these areas in the decision-making process. To date, however, a causal test of this role is lacking.

Here, we investigated whether M1 causally contributes to deliberation about action choices, or whether it simply reflects decision signals coming from upstream areas, such as PMd, the basal ganglia, or the locus coereleus. It has previously been shown that disrupting M1 activity by means of continuous theta burst stimulation (cTBS) causes finger responses to slow down (Huang et al., 2005; Lakhani et al., 2014; McAllister et al., 2013). However, it is a matter of debate whether this effect should be interpreted as a slowing down of processes involved in deciding which action to perform (*i.e.*, in the deliberation process) or as a slowing down of the motor processes that follow commitment (*i.e.*, of movement initiation and execution). In fact, a slowing down of the deliberation process has been associated with reduced urgency during volitional decision behavior (Hanks et al., 2011; Seideman et al., 2018; Thura and Cisek, 2014a, 2016). If M1 is causally involved in the decision process, then M1 disruption might lengthen deliberation in a manner consistent with reduced urgency as compared to a sham cTBS session. Conversely, if M1 is mainly involved in initiating and executing selected actions, then its disruption should have no effect on the deliberation portion of response time, and should only slow down the motor processes that follow commitment to an action. These hypotheses were tested by characterizing action choices in a modified version of the tokens task (Cisek et al., 2009), which is specifically designed to estimate subjects’ time of commitment, their accuracy criterion and infer from those variables their urgency functions (Thura et al., 2014a).

## MATERIALS AND METHODS

### Participants

19 healthy right-handed subjects participated in this study (10 women; 24 ± 3.5 years old). Participants were financially compensated for their participation and earned additional money depending on their performance in a decision-making task (see *Task* section below). The protocol was approved by the institutional review board of the catholic University of Louvain (UCLouvain), Brussels, Belgium, and required written informed consent, in compliance with the principles of the Declaration of Helsinki.

### Experimental design

Experiments were conducted in a quiet and dimly-lit room. Subjects were seated at a table in front of a 21-inch cathode ray tube computer screen. The display was gamma-corrected and its refresh rate was set at 100 Hz. The computer screen was positioned at a distance of 70 cm from the subject’s eyes and was used to display stimuli during the decision-making task. Left and right forearms were rested upon the surface of the table with the palms facing the table. A computer keyboard was positioned upside-down under the dominant (*i.e.*, right) hand with the response keys F9 and F8 under the index and middle fingers, respectively (see Figure 1).

**Fig.1.**
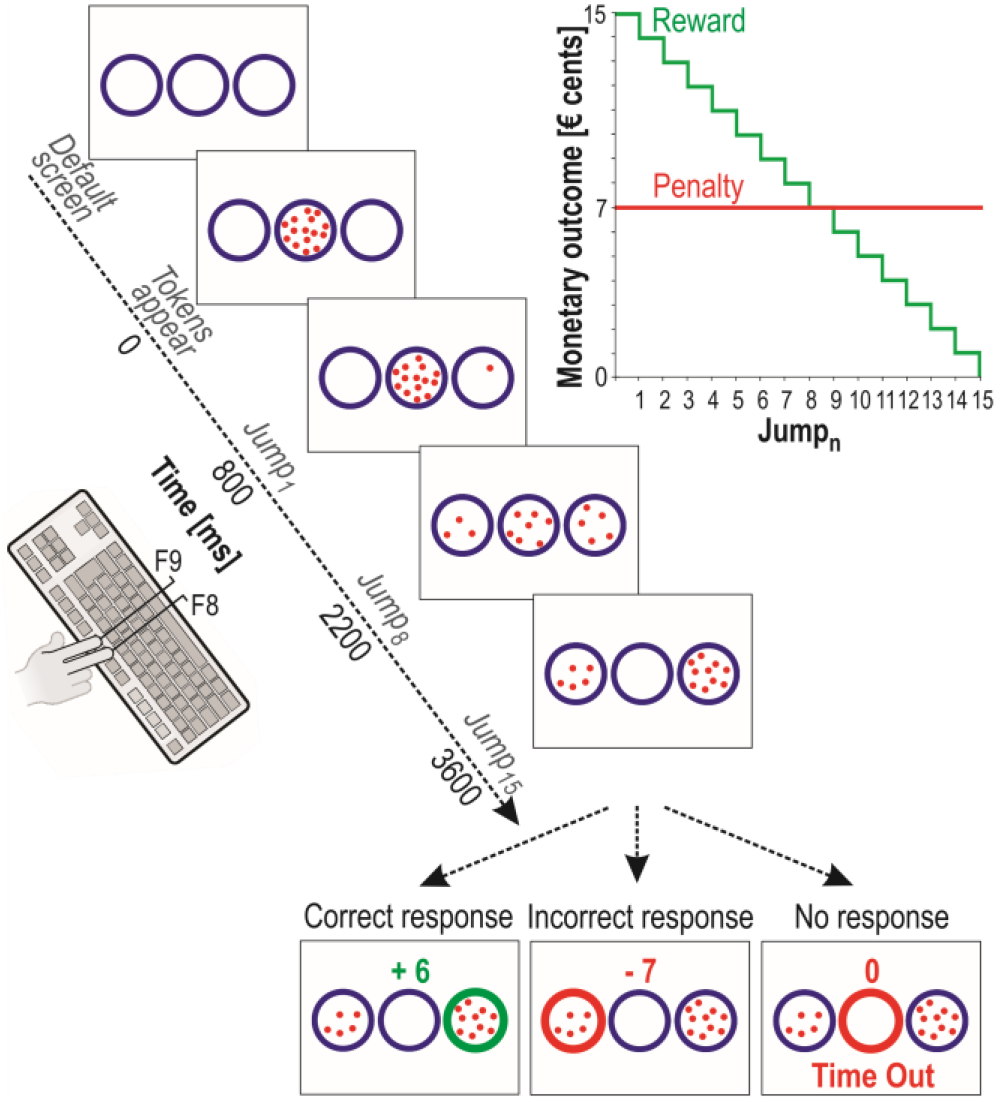
Schematic of the tokens task. In each trial, 15 tokens jumped one-by-one every 200 ms from the central circle to one of the two lateral circles (i.e., Jump_1_ to Jump_15_). The subjects had to indicate by a right index or right middle finger key-press (i.e., F9 and F8 keys, respectively) which lateral circle they thought would receive more tokens (i.e., left or right circle, respectively) at the end of the trial. They could provide their response whenever they wanted between Jump_1_ and Jump_15_. For a correct response, the subjects earned, in € cents, the number of tokens remaining in the central circle at the time of the response. Hence, the reward received for a correct response decreased over time, as depicted on the upper right side of the figure (green trace). The example presented on the lower left side of the figure represents a correct response provided between Jump_9_ and Jump_10_ (i.e., the score indicates that 6 tokens remained in the central circle at the moment the right circle was chosen). In contrast, if subjects chose the incorrect lateral circle, they lost 7 € cents, regardless of their RT. As such, the penalty score was fixed, as shown in red on the upper right side of the figure: the lower middle example represents an incorrect choice of the left circle. Thus, the reward/penalty ratio decreased over time, producing an increasing sense of urgency over the course of a trial. In the absence of response (“Time Out” trial, lower right side example), subjects were neither rewarded, nor penalized (score = 0). For representative purposes, the “Time Out” message is depicted below the circles in this example, while it was presented above them in the experiment.

### Task

The task used in the current study is a variant of the “tokens task” (Cisek et al., 2009) and was implemented by means of LabView 8.2 (National Instruments, Austin, TX). The sequence of events in each trial is depicted in Figure 1. Between trials, subjects were always presented with a default screen consisting of three circles (4.5 cm diameter), displayed for 2000 ms on a white background. Fifteen randomly arranged tokens (0.3 cm diameter) then appeared in the central circle. After a delay of 800 ms, the tokens began to jump, one-by-one every 200 ms from the center to one of the two lateral circles (*i.e.*, Jump_1_ to Jump_15_). The subjects’ task was to indicate by a right index or right middle finger key-press which lateral circle they thought would ultimately receive the majority of the tokens (*i.e.*, F9 and F8 key-presses to choose left and right circles, respectively). They could provide their response as soon as they felt sufficiently confident, but between Jump_1_ and Jump_15_. Once the response was provided, the tokens kept on jumping every 200 ms until the central circle was empty. At this time, the selected circle was highlighted either in green or in red depending on whether the response was correct or incorrect, respectively, and a score was displayed above the central circle to provide the subjects with further feedback of their performance. In correct trials, subjects received a positive score (*i.e.*, a monetary reward) which was equal to the number of tokens remaining in the central circle at the time of the response (in € cents). Conversely, incorrect responses led to a fixed penalty of 7 cents, regardless of the RT. Thus, the longer the subjects waited to provide a response, the lower was the reward/penalty ratio, generating an increasing sense of urgency as time passed within each trial. In the absence of any response before Jump_15_, the central circle was highlighted in red and a “Time Out” (TO) message appeared on the top of the screen. The subjects were neither rewarded nor penalized in these trials. The feedback cue remained on the screen for 1000 ms and then disappeared at the same time as the tokens did, denoting the end of the trial. Subjects were told that they would receive a monetary reward at the end of the experiment corresponding to their final score. Each trial lasted 6600 ms.

### Blocks and sessions

The study included 3 sessions, conducted on separate days at a 24-hour interval. Testing always occurred at the same time of the day for a given subject, to avoid variations that could be due to changes in chronobiologic states (Derosiere et al., 2015a; Schmidt et al., 2006). Each session comprised 4 blocks of 50 trials, with each block lasting about 5.5 minutes. Subjects also performed 4 blocks of 5 trials of a simple reaction time (SRT) task, two at the beginning and two at the end of each session. In the SRT task, subjects were presented with the same display as in the tokens task described above. However, after 50 ms in the central circle, the 15 tokens all jumped together into one of the two lateral circles at the same time. Subjects were instructed to respond to this “GO signal” by pressing the corresponding key as fast as possible. Importantly, the 15 tokens always jumped into the same lateral circle in all trials of a given SRT block and subjects were told which lateral circle this would be in advance. This SRT task allowed us to estimate the sum of the delays attributable to the sensory and motor processes in the absence of a choice (see Cisek et al., 2009; Thura et al., 2014).

Day 1 served as a training session. Day 2 and 3 corresponded to the actual experimental sessions with the cTBS intervention (Figure 2). cTBS was applied before subjects engaged in the blocks of trials, either over the left M1 hand area (M1-Disruption session) or over the right primary somatosensory cortex (S1), 2 cm behind the right M1 area (Sham session; Derosiere et al., 2014; Alexandre et al., 2015; Torta et al., 2013), in a randomized order. The Sham session allowed us to ensure that the putative behavioral effects observed following M1 cTBS were not due to the tactile and auditory sensations elicited by the TMS pulses (Derosiere et al., 2017a, 2017b).

**Fig. 2.**
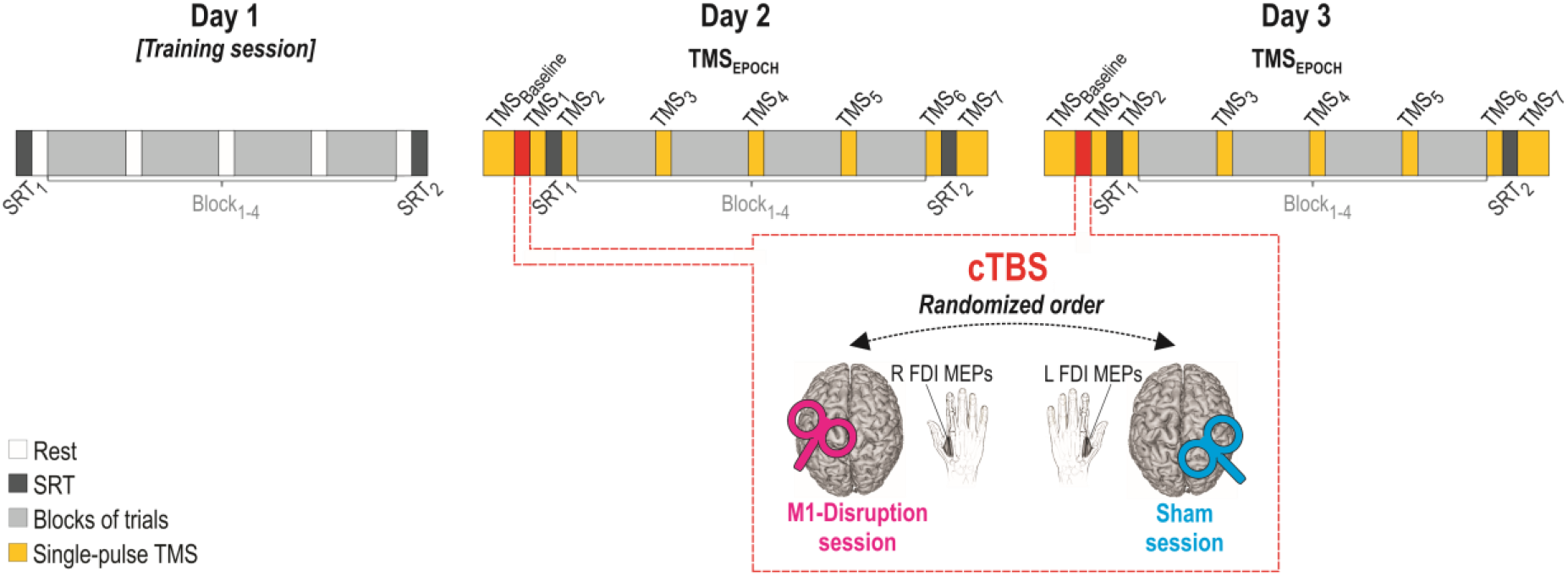
Experimental protocol. Subjects came to the lab for 3 consecutive days (Day 1, 2 and 3). On each day, they performed the tokens task during 4 blocks of 50 trials (Block_1-4_; light grey rectangles). They also performed Simple Reaction Time blocks at the beginning and at the end of each session (SRT_1_ and SRT_2_, respectively; dark grey rectangles). Day 1 served as a training session and did not involve any continuous Theta Burst Stimulation (cTBS). A cTBS train was applied for 40 s at the beginning of Day 2 and 3 (red rectangles), either over the left (L) primary motor cortex (M1-Disruption session, magenta coil) or over the right (R) primary somatosensory cortex (Sham session, cyan coil), in a randomized order. Motor Evoked Potentials (MEPs) were elicited at different time points (TMS_EPOCHS_) throughout the sessions (TMS_Baseline_ and TMS_1-7_, yellow rectangles), either in the R first dorsal interosseous (FDI) muscle (M1-Disruption session) or in the L FDI (Sham session), by applying single-pulse TMS over the L or R M1, respectively. Note that in the Sham session, this implied targeting different sites for the cTBS intervention (R S1) and the MEP assessments (R M1; coil position not shown on the figure).

### TMS procedure

TMS was delivered through a 2×75 mm figure-of-eight coil connected to a Magpro X100 Stimulator (Magventure Company, Farum, Denmark). The coil was placed tangentially on the scalp with the handle oriented towards the back of the head and laterally at a 45° angle away from the midline. At the beginning of each session, the M1 hand area was localized by identifying the optimal spot (called the “hotspot”) for eliciting MEPs in the first dorsal interosseous (FDI) muscle of the right hand (M1-Disruption session) or the left hand (Sham session). To do so, we relied on markers disposed on an electroencephalography (EEG) cap fitted on the participant’s head (Duque et al., 2010, 2014; Vandermeeren et al., 2009). We first applied the stimulation with the center of the coil over the C3 or C4 location of the EEG cap (*i.e.*, corresponding to the right and left M1 areas, respectively). Stimulation intensity was increased until obtaining consistent MEP responses at this location. We then moved the coil by steps of approximately 0.5 cm around this location both in the rostro-caudal and in the medio-lateral axis. Stimulation was applied with the previously defined intensity at each new location and MEP amplitudes were visually screened. The hotspot was defined as the location at which the largest and most consistent MEP amplitudes could be obtained. The coil was then held at this location and the edges of its shape were marked on tapes disposed on the EEG cap. These marks allowed us to localize the hotspot at any required time during the session. Once the hotspot was found, we determined the resting motor threshold (rMT). The rMT was defined as the minimal intensity required to evoke MEPs of 50 μV peak-to-peak in the targeted muscle on 5 out of 10 consecutive trials at the hotspot (Grandjean et al., 2018; Rothwell et al., 1991; Rossini et al., 1994, 2015; Vassiliadis et al., 2018).

The cTBS procedure consisted of a series of short TMS trains (three pulses at 50 Hz) repeated every 200 ms for a total duration of 40 s (600 pulses) at an intensity of 80 % of rMT (Derosiere et al., 2017a, 2017b; Huang et al., 2005; Solopchuk et al., 2017). Such an intervention has been shown to inhibit the stimulated cortical area, producing a temporary “virtual lesion”, often effective as soon as the train is over (Derosiere et al 2017a, 2017b; Do et al., 2018; Sasaki et al. 2018) and lasting for between 20 (Clerget et al., 2011; Oberman et al., 2011; Zénon et al., 2015) and 45 minutes (Huang et al., 2005).

In order to monitor the inhibitory effect of cTBS on motor activity, single TMS pulses were applied at the M1 hotspot at 115 % of the rMT to elicit MEPs at different time points in the M1-Disruption and Sham sessions (Klein et al., 2014; Labruna et al., 2014; Quoilin et al., 2016, 2017). In the M1-Disruption session, MEPs were recorded in the right FDI following TMS over left M1, to evaluate the impact of left M1 cTBS on left motor excitability. In the Sham session, MEPs were obtained from the left FDI following TMS over right M1, to control for the absence of effect of right S1 cTBS on right M1 excitability.

The time points at which MEPs were elicited in the M1-Disruption and Sham sessions were comparable (see Figure 2). In both sessions, 20 MEPs were elicited at the beginning of the session (*i.e*., just before cTBS; TMS_Baseline_). Then, 15 MEPs were elicited just following cTBS (TMS_1_), after the two initial SRT blocks (TMS_2_), and after each block of trials of the tokens task (TMS_3-6_). Finally, 20 additional MEPs were evoked following the two last SRT blocks (TMS_7_). These seven timings (TMS_1-7_) fell 1 min, 3 min, 11 min, 19 min, 27 min, 35 min, and 37 min after the cTBS intervention, respectively.

### Data collection

Electromyography (EMG) was used to measure the peak-to-peak amplitude of FDI MEPs elicited by single TMS pulses over the contralateral M1. EMG activity was recorded from surface electrodes placed over the right FDI or the left FDI (M1-Disruption or Sham sessions, respectively). EMG data were collected for 1000 ms on each trial, starting 300 ms before the TMS pulse. EMG signals were amplified, bandpass filtered on-line (10-500 Hz) and digitized at 2000 Hz for off-line analysis.

### Data analysis

#### Motor evoked potential data

MEP data were collected with Signal (Signal 3.0, Cambridge, UK) and analyzed with custom Signal scripts. MEP amplitudes were measured for each TMS pulse. Trials with background EMG activity greater than 20 μV on average (root mean square, rms), in the 200-ms window preceding the TMS artifact, were excluded from the analysis. 3.63 ± 5.43 % of trials were discarded based on this criterion. The amplitude of MEPs elicited at each time point was averaged to obtain a measure of motor excitability at TMS_Baseline_ and at TMS_1-7_ in the M1-Disruption and Sham sessions (Figure 2). For each session, we then expressed MEP amplitudes obtained at TMS_1-7_ (*i.e.*, after the cTBS intervention) in percentage of the amplitudes measured at TMS_Baseline_ (*i.e.*, before the cTBS intervention).

The disruptive impact of cTBS on cortical activity varies between subjects (Do et al., 2018; Jannati et al., 2017; Rocchi et al., 2018). Here, we aimed at only including individuals in which cTBS effectively disrupted M1. To do so, we discarded subjects presenting percentage MEP amplitudes exceeding 2.5 SD *above* the mean of the group in the M1-disruption session at one of the TMS_EPOCHS_ or more. This led to the rejection of three subjects, who exhibited average MEP amplitudes of 148.3 ± 12.5 %, 139.5 ± 10.6 % and 188.6 ± 20.9% following the cTBS intervention in the M1-disruption session (all TMS_EPOCHS_ averaged together), reflecting thus a large increase (rather than the targeted decrease) in motor excitability. The analyses of the MEP and behavioral data were performed on the remaining pool of subjects (n = 16).

#### Behavioral data

Behavioral data were collected with LabVIEW 8.2 (National Instruments, Austin, TX), stored in a database (Microsoft SQL Server 2005, Redmond, WA), and analyzed with custom MATLAB scripts (MathWorks, Natick, MA). Because Day 1 served as a training session (see section *Blocks and sessions*), the behavioral analyses focused on the data acquired on Day 2 and 3 (*i.e*., in the M1-Disruption and Sham sessions).

#### SRT task

For each subject, we computed the mean SRT for both fingers (right index and right middle fingers), defined as the difference between the time at which subjects pressed the key and the time at which the 15 tokens appeared simultaneously in the lateral circle, obtained at the beginning (SRT_1_) and at the end (SRT_2_) of each session. This SRT allowed us to quantify the impact of M1 cTBS on the motor processes that follow commitment to an action in the absence of a choice.

#### Tokens task

##### Classification of the trial types based on the temporal profile of the success probability

The task allows us to calculate, at each moment in time, the “success probability” *p*_*i*_*(t)* associated with choosing each lateral circle *i*. For a total of 15 tokens, if at a particular moment in time the right (R) circle contains N_R_ tokens, the left (L) circle contains N_L_ tokens, and the central (C) circle contains N_C_ tokens, then the probability that the circle on the right will ultimately be the correct one is described as follows:

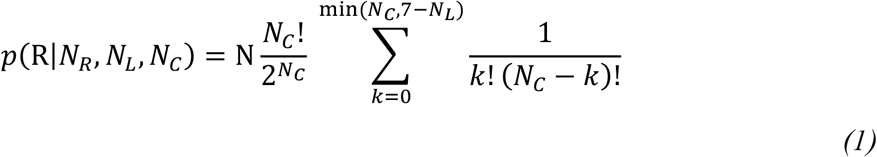

For some of the analyses, we grouped trials according to the temporal profile of *p*_*i*_*(t)*. That is, although the side of each token jump was completely random in each trial, we could classify some trials as belonging to one of two specific types a posteriori. Trials were categorized as “obvious” when the *p*_*i*_*(t)* was above 0.6 after Jump_2_ and above 0.75 after Jump_5_; that is, the initial token jumps consistently favored the correct circle. Other trials were categorized as “ambiguous” when the initial jumps were balanced between the lateral circles, keeping the *p*_*i*_*(t)* close to 0.5 until late in the trial: *p*_*i*_*(t)* remained between 0.5 and 0.66 up to Jump_7_ in these trials.

##### Decision Time (DT), percentage of correct choices (%Correct) and percentage of time out trials (%TO)

For each session (M1-Disruption and Sham) and each trial type (obvious and ambiguous; trials that were neither obvious nor ambiguous were not considered here), we analyzed the following behavioral variables: the decision time (DT), the percentage of correct choices (%Correct) and the percentage of time out trials (%TO). To evaluate the DT, we first calculated the RT during the tokens task by computing the difference between the time at which subjects pressed the key and Jump_1_. We then subtracted from this tokens RT, the mean RT obtained in the SRT task on the same day (SRT_1_ and SRT_2_ pooled together), providing us with an estimate of DT, reflecting the duration of the deliberation process for each subject. Note that we used a monetary reward in the tokens but not in the SRT task, which might have led us to slightly underestimate the DT. That is, previous studies have shown that monetary reward can boost motor processes (Reppert et al., 2018; Summerside et al., 2018; Yoon et al., 2018). Hence, the latter might have been faster in the tokens than in the SRT task used to estimate it. Thus, we might have subtracted a too large value from the tokens RT, shortening the DT. Still, this putative underestimation of DT applies for both the M1-disruption and the Sham sessions and has thus no biasing impact on our data.

##### Sensory evidence at decision time (DT)

Sensory evidence refers to the available information supporting the correct choice. In the tokens task, the sensory evidence is determined by the difference between the number of tokens in each lateral circle; the more the correct circle contains a large number of tokens, compared to the other lateral circle, the higher the evidence. Given that tokens jump one by one in this task, sensory evidence changes after each jump. We can estimate the evidence based on which subjects made their decision by computing after each jump a first-order approximation of the real probability function (equation 1), called the sum of log-likelihood ratios (SumLogLR), and then compute this quantity at decision time (Cisek et al., 2009):

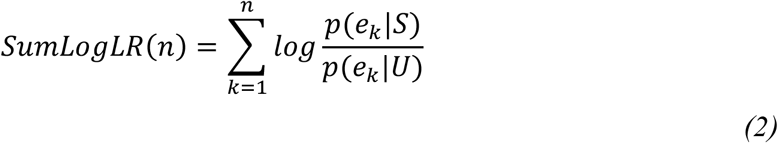

In this equation, *p(e*_*k*_|*S)* is the likelihood of a token event e_k_ (a token jumping into either the selected or unselected lateral circle) during trials in which the selected lateral circle *S* is correct and *p(e*_*k*_|*U)* is the likelihood of e_k_ during trials in which the unselected circle *U* is correct. The SumLogLR is proportional to the difference between the number of tokens that moved towards each lateral circle before the decision. Hence, the lower the amount of sensory evidence in favor of the chosen lateral circle, the lower the SumLogLR.

To characterize the decision policy of the subjects in the Sham and M1-Disruption sessions, we determined the level of sensory evidence at the time of commitment (*i.e.*, at DT). To do so, we binned trials as a function of the total number of tokens that moved before the decision, and calculated the average SumLogLR for each bin as performed in previous studies exploiting the tokens task (*e.g.*, Thura and Cisek, 2014, 2017). Seven bins were defined, with the first bin (Bin_1_) including responses provided between Jump_5_ and Jump_6,_ the second bin (Bin_2_) including responses provided between Jump_6_ and Jump_7_ and so on, until the last bin (Bin_7_) covering the period between Jump_11_ and Jump_12_. SumLogLR at DT preceding Jump_5_ or following Jump_12_ were not considered for this analysis because part of the subjects did not respond at these timings. Importantly, the SumLogLR at DT was computed based on every trial where a response was provided (*i.e.*, for correct and incorrect responses, in obvious and ambiguous trials, as well as in other trials with different *p*_*i*_*(t)*).

##### Estimation of urgency functions

According to recent models of decision-making, action choices result from the combination of signals that track the available sensory evidence and the level of urgency that grows over time (Cisek et al., 2009, Ditterich, 2006, Drugowitsch et al., 2012). For instance, in a minimal implementation of the urgency gating model (Cisek et al., 2009; Thura et al., 2012), evidence is multiplied by a linearly increasing urgency signal, and then compared with a threshold. The result can be expressed as follows:

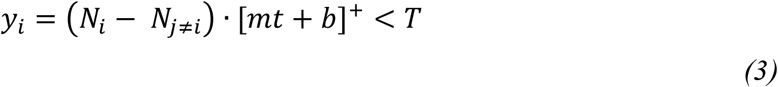

where *y*_*i*_ is the “neural activity” for choices to target *i*, *N*_*i*_ is the number of tokens in target *i*, *t* is the number of seconds elapsed since the start of the trial, *m* and *b* are the slope and y-intercept of the urgency signal, and []^+^ denotes half-wave rectification (which sets all negative values to zero). When *y*_*i*_ for any target crosses the threshold *T*, that target is chosen.

A direct prediction of such urgency-based models is that decisions made with low levels of evidence should be associated with high levels of urgency, and vice-versa. That is, one core assumption is that a high urgency should push one to commit to a choice even if evidence for that choice is weak. Hence, the SumLogLR at DT values (*i.e.*, reflecting the available sensory evidence at the time of commitment in the tokens task) can be exploited to estimate the level of urgency at DT. Here, we first multiplied the SumLogLR at DT values by −1 (*i.e.*, to “rectify” them; please see Figure 3), given the theoretical inverse relationship between sensory evidence and urgency at DT. We then added a constant of 3 to the rectified curves to obtain positive values. Finally, we fitted a linear regression over the rectified positive values. We extracted the intercept and the slope of these so-called urgency functions, which we used as estimates of the initial level and the growth rate of the urgency signal, respectively. Second-order polynomial regressions were also performed on the rectified SumLogLR data but did not yield significantly better fits (*i.e.*, group-level Bayesian Information Criterion [BIC] values for linear and polynomial fits were 5.75 ± 1.61 and 5.74 ± 1.79 a.u., respectively; t_14_ = 0.02, p =.984).

**Fig.3.**
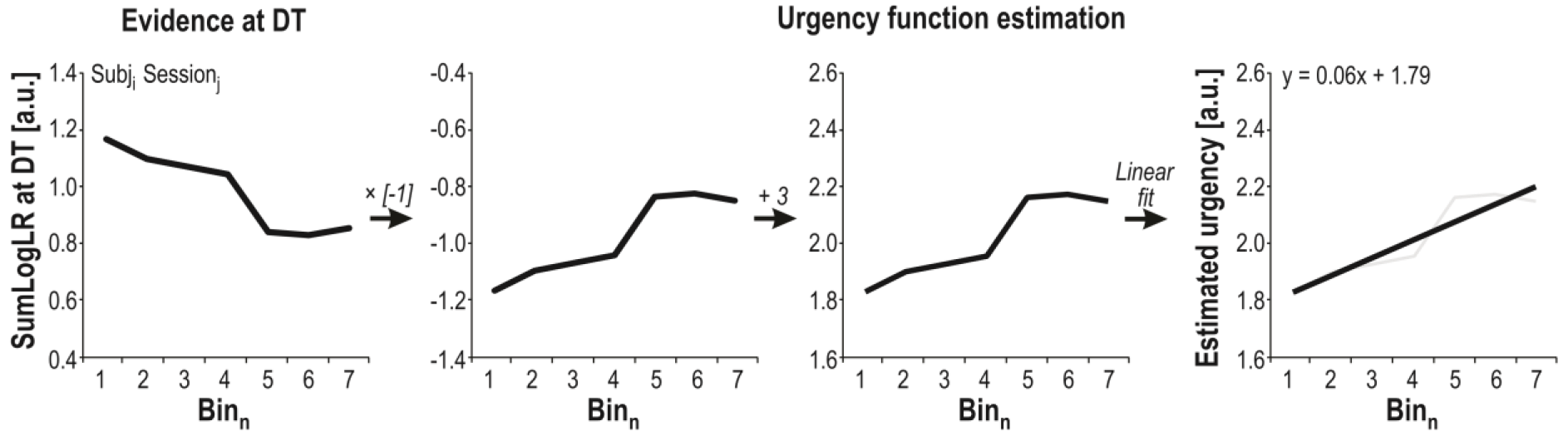
Urgency function estimation. We exploited the SumLogLR values to estimate the level of urgency at DT for each subject *i* and each session *j*. To do so, we followed four steps (shown from left to right). First, SumLogLR values were obtained for different bins of DT. Then, these values were multiplied by −1. Next, a constant of 3 was added to obtain positive values. Finally, a linear regression was fitted over the positive values. The equation of the regression allowed us to extract the intercept and the slope of the obtained urgency function (1.79 and 0.06 in this example, respectively).

### Statistical analysis

All statistical analyses were performed with custom R Scripts (R Version 3.4.1, Car and BayesFactor packages). All data were examined for normality and homogeneity of variance using Skewness, Kurtosis and Brown-Forsythe tests. The significance level for all tests was set at p < .05, except when Bonferroni corrections were applied. All results are expressed as mean ± SE.

#### Motor evoked potential data

MEP data (expressed in percentage of MEPs at TMS_Baseline_) were analyzed using a two-way repeated-measure ANOVA (ANOVA_RM_) with SESSION (M1-Disruption, Sham) and TMS_EPOCH_ (TMS_1_, TMS_2_, TMS_3_, TMS_4_, TMS_5_, TMS_6_, TMS_7_) as within-subject factors. Moreover, the percentage MEP values obtained for each TMS_EPOCH_ were compared against 100 % using Bonferroni-corrected single-sample Student’s t-tests to identify any significant suppression in the M1-Disruption and in the Sham session.

#### Behavioral data

The SRT data were analyzed using a two-way ANOVA_RM_ with SESSION (M1-Disruption, Sham) and SRT_EPOCH_ (SRT_1_, SRT_2_) as within-subject factors. The DT, the %Correct and the %TO data were analyzed using two-way ANOVA_RM_ with SESSION (M1-Disruption, Sham) and TRIAL (obvious, ambiguous) as within-subject factors. The SumLogLR at DT was analyzed using a two-way ANOVA_RM_ with SESSION (M1-Disruption, Sham) and BIN (Bin_1_ to Bin_7_) as within-subject factors. When appropriate, Tukey’s HSD post-hoc tests were used to detect paired differences in these ANOVAs. Furthermore, the intercept and the slope of the urgency functions were compared between the two sessions using Student’s t-tests.

## RESULTS

### Motor evoked potential data

The ANOVA_RM_ revealed a main effect of the factor SESSION on the percentage MEP amplitudes (F_1,15_ = 15.41; p = .001; see Figure 4). As such, percentage MEP amplitudes were lower following cTBS in the M1-Disruption session (78.78 ± 3.04 %) than in the Sham session (105.53 ± 5.32 %; TMS_1-7_ timings pooled together). The effect size (Cohen’s d) for this factor was 1.5 indicating a large effect of SESSION (Cohen, 1988). This effect on percentage MEP amplitudes did not depend on time as the ANOVA_RM_ did not reveal any effect of the factor TMS_EPOCH_ (F_6,90_ = 1.99, p = .075) nor interaction with the factor SESSION (F_6,90_ = 0.72, p = .637). Hence, percentage MEP amplitudes remained stable following the cTBS intervention; they were consistently lower in the M1-Disruption than in the Sham session, regardless of the time at which MEPs were considered during the course of the experiment.

**Fig. 4.**
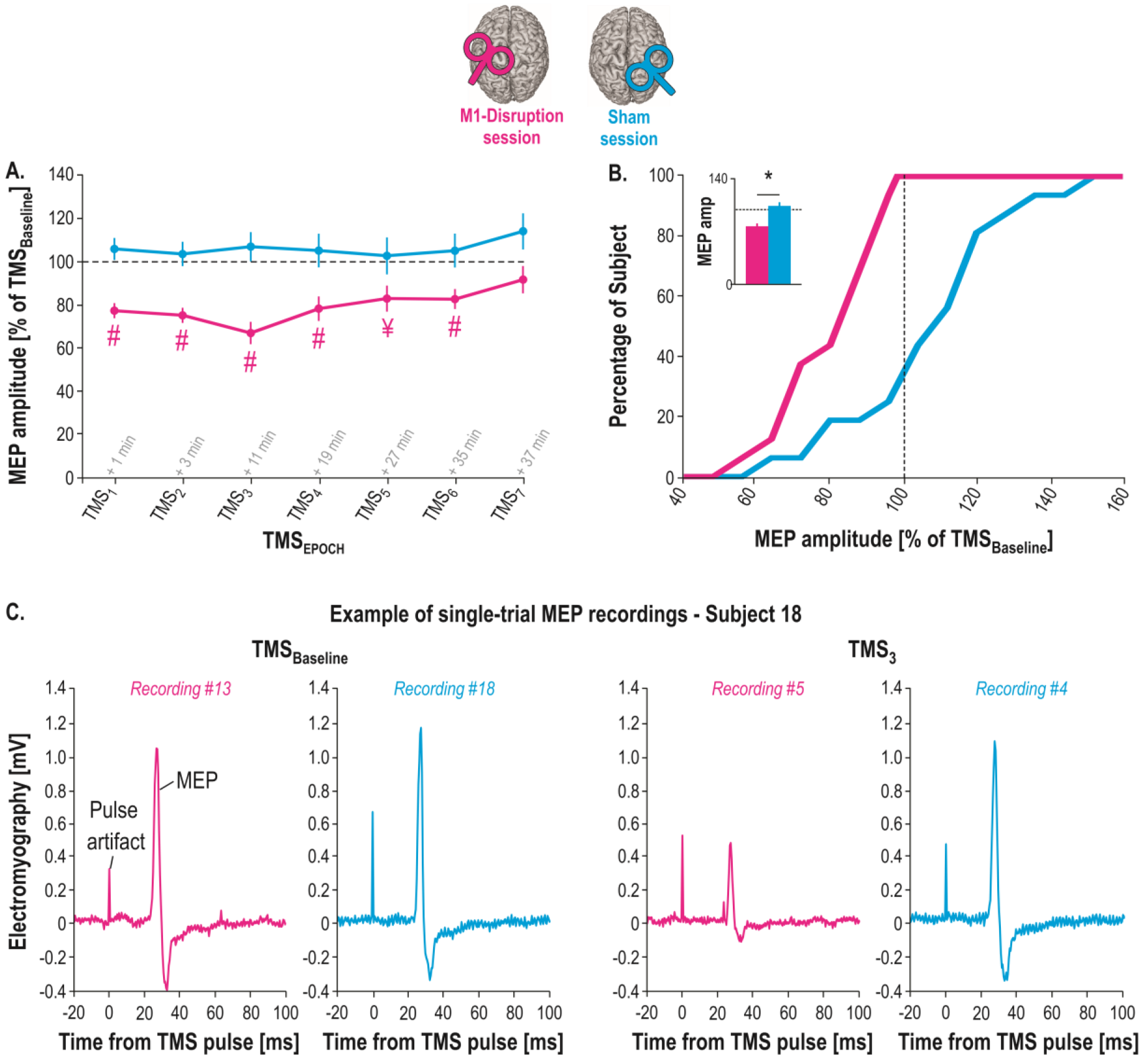
Motor Evoked Potentials (MEPs) amplitudes. *A.* Mean value of MEPs (in percentage of MEPs at TMS_Baseline_) elicited after the cTBS intervention in the First Dorsal Interosseous (FDI) muscle at each of the TMS_EPOCHS_ (TMS_1-7_) in the M1-Disruption (magenta traces) and Sham (cyan traces) sessions. Note the significant disruption of MEPs with respect to baseline (*i.e.*, dashed horizontal line) in the M1-Disruption session (#: significantly different from 100 at p < .0036 [Bonferroni-corrected]; ¥: significantly different from 100 at p < .05 [uncorrected; p = .017]). *B.* Cumulative percentage of subjects. MEPs obtained at TMS_1-7_ are pooled together. Note that all subjects included in the analysis showed percentage MEP amplitudes smaller than 100% in the M1-disruption (*i.e.*, a disruptive effect), while the same subjects did not show any effect in the Sham session, as also shown in the inset representing the group-level average with the effect of SESSION (*: significantly different at p < .05). Error bars represent SE. *C.* Example of single-trial MEP recordings Each trace depicts a raw EMG signal in a representative subject (#18), starting 20 ms before the TMS pulse and ending 100 ms after it. The artifact caused by the pulse is reflected as a peak occurring at time 0; the MEP occurs approximately 22 ms later. The four recordings display MEPs elicited at TMS_Baseline_ (left) or TMS_3_ (right) in the M1-Disruption (magenta traces) or Sham (cyan traces) session. In this subject, cTBS had a strong effect; average percentage MEP amplitudes at TMS_3_ were much smaller in the M1-Disruption session (66 ± 12.8 %) compared to the sham session (112.6 ± 5.7 %).

Additional single-sample Student’s t-tests against 100 % (run for each TMS_EPOCHS_; Bonferroni-corrected at p < .0035) showed that, as expected, the difference in MEP amplitude between the two sessions reported above was due to a selective suppression of MEPs in the M1-Disruption but not in the Sham session. As such, percentage MEP amplitudes were significantly lower than 100 % at almost all timings in the M1-Disruption session, except for TMS_5_ and TMS_7_ (*i.e.*, at TMS_1-4_ and TMS_6_; all p-values = [.000008 .002]). Conversely, amplitudes were never significantly different from 100 % (*i.e.*, from TMS_Baseline_) in the Sham session (all p-values = [.163 .826]), indicating that right S1 cTBS had no impact on right M1 activity, as previously reported (Derosiere et al., 2017a, 2017b).

### Behavioral data

#### SRT Task

The ANOVA_RM_ revealed a main effect of the factor SESSION on the SRT data (F_1,15_ = 5.34, p = .035; see Figure 5.A). Indeed, SRTs were significantly prolonged in the M1-Disruption session (237.3 ± 7.4 ms) compared to the Sham session (220.8 ± 5.6 ms; SRT_1_ and SRT_2_ pooled together). The effect size (Cohen’s d) for this factor was 0.6 indicating a medium to large effect of SESSION. The impact of M1 disruption on SRTs did not vary over the course of the session. As such, the ANOVA_RM_ did not reveal any significant effect of the factor SRT_EPOCH_ (F_1,15_ = 0.02, p = .892) nor of its interaction with the factor SESSION (F_1,15_ = 1.78, p = .202). These findings indicated that M1-disruption altered the motor processes underlying initiation and/or execution of the cued movements (Huang et al., 2005; Lakhani et al., 2014; McAllister et al., 2013).

**Fig. 5.**
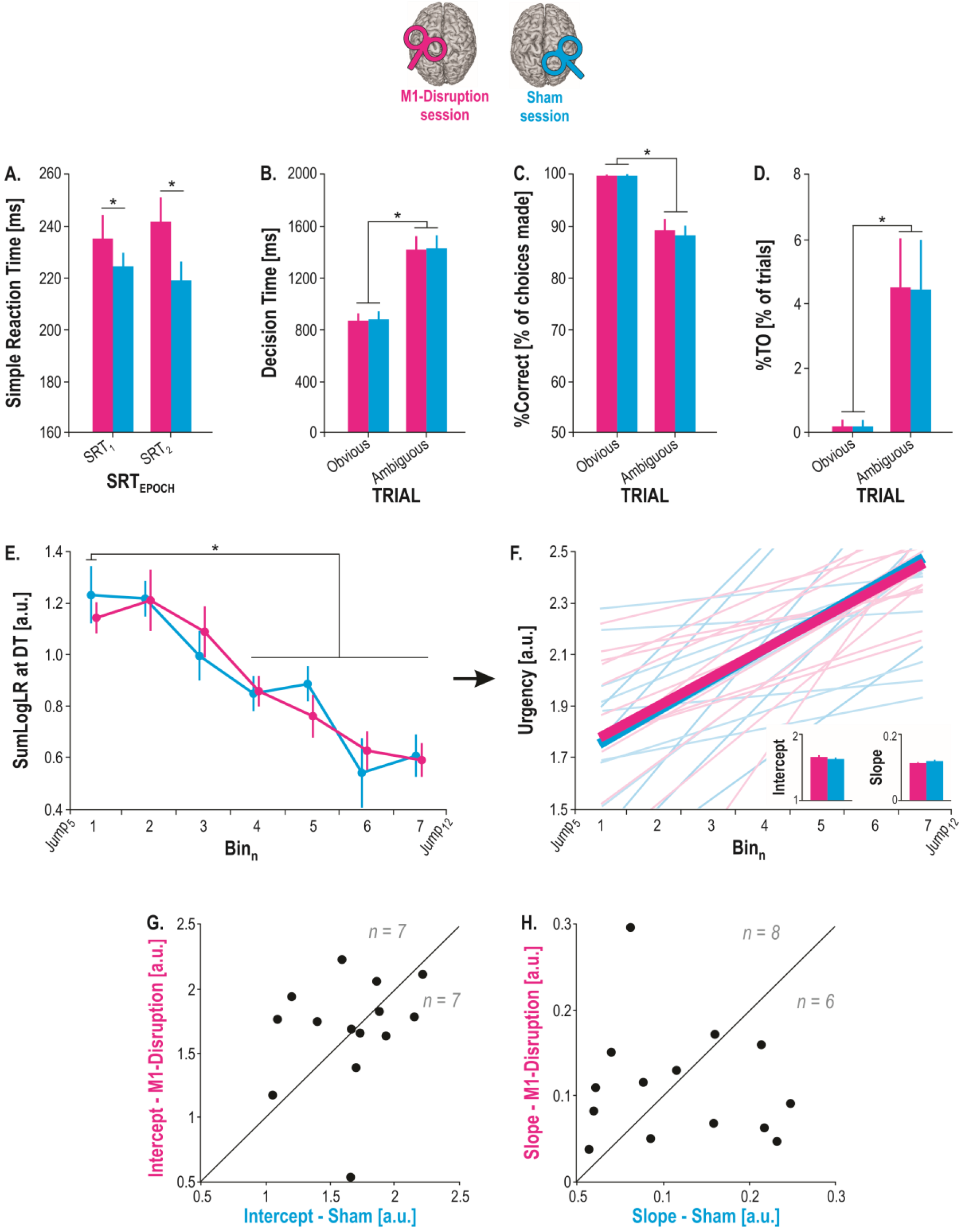
Behavioral results. *A.* Mean Simple Reaction Time (SRT) (obtained in the SRT task) measured at each of the SRT_EPOCHS_ (SRT_1-2_) in the M1-Disruption (magenta bars) and Sham (cyan bars) sessions. *B.* Mean Decision Time (DT) measured in each TRIAL (obvious, ambiguous) in the M1-Disruption (magenta bars) and Sham (cyan bars) sessions. *C* and *D* same as B for the percentages of correct choices (%Correct) and of Time Out trials (%TO), respectively. *E.* SumLogLR at DT measured in each Bin_n_ of decision time (*i.e.*, between Jump_5_ and Jump_12_, see main text) in the M1-Disruption (magenta traces) and Sham (cyan traces) sessions. *F.* Urgency functions computed based on the rectified SumLogLR at DT for the M1-Disruption (magenta traces) and Sham (cyan traces) sessions. The small bar graphs on the bottom right represent the group-level average intercept and slope of the functions. Light lines illustrate individual estimated urgency functions, bold lines illustrate the mean urgency functions averaged across population. *G.* Individual intercept values, represented for the M1-Disruption Session (y-axis) as a function of the values for the Sham session (x-axis). Points above the diagonal (n = 7/14) represent the subjects showing a higher intercept in the M1-Disruption than following Sham session, while points below the diagonal (n = 7/14) represent the subjects showing a lower intercept in the M1-Disruption than following Sham session. *H.* Same as G. for the slope values. *: significant difference (p < .05). Error bars represent SE.

#### Tokens task

##### Decision time (DT)

As expected, the ANOVA_RM_ revealed a main effect of the factor TRIAL on the DT data (F_1,15_ = 119.30, p < .00001; see Figure 5.B). Indeed, DTs were significantly shorter in obvious trials (871.77 ± 57.14 ms) than in ambiguous ones (1426.57 ± 88.59 ms; M1-Disruption and Sham sessions pooled together). Importantly, the ANOVA_RM_ did not reveal any significant effect of the factor SESSION (F_1,15_ = 0.24, p = .631) nor of its interaction with the factor TRIAL (F_1,15_ = 0.02, p = .888). Hence, the time taken by the subjects to deliberate depended on the trial type they encountered (*i.e.*, obvious vs ambiguous) but was not affected by M1 disruption.

##### Percentage of correct choices (%Correct)

We found a significant main effect of the factor TRIAL for the %Correct data (F_1,15_ = 31.727, p = .00005; see Figure 5.C). Indeed, %Correct was significantly higher in obvious trials (99.80 ± 0.13 %) than in ambiguous ones (88.61 ± 1.80 %; M1-Disruption and Sham sessions pooled together). However, neither the factor SESSION (F_1,15_ = 0.05, p = .497) nor its interaction with the factor TRIAL were significant (F_1,15_ = 0.44, p = .517). Hence, M1 disruption did not alter the accuracy of the decision process.

##### Percentage of Time Out trials (%TO)

The %TO data revealed a similar pattern as the variables described above. Indeed, the %TO was significantly lower in obvious (0.20 ± 0.13 %) than in ambiguous trials (4.43 ± 1.42 %; M1-Disruption and Sham sessions pooled together), as confirmed by the ANOVA_RM_ (factor TRIAL: F_1,15_ = 7.47, p = .015; see Figure 5.D). Moreover, there was no effect of the factor SESSION (F_1,15_ = 0.01, p = .920) or interaction with the factor TRIAL (F_1,15_ = 0.01, p = .922). Hence, the proportion of trials in which subjects refrained from responding was unaffected by M1-Disruption.

##### Sensory evidence at decision time (SumLogLR at DT) and urgency

The amount of sensory evidence based on which subjects made their decision was estimated using the SumLogLR (computed at DT): the higher the SumlogLR, the higher the evidence at DT (Cisek, 2009; Thura et al., 2012, 2014). SumLogLR values are presented for each Bin_n_ on Figure 5.E (see *Methods*), separately for the M1-Disruption and the Sham sessions. Note that two subjects were excluded from this analysis as they responded too early on most trials, resulting in a lack of SumLogLR values after Jump_9_ in these participants. Hence, SumLogLR analyses were run on 14 subjects.

Overall, fast decisions were made based on more sensory evidence than slow decisions, as confirmed by the ANOVA_RM_ showing a main effect of the factor BIN on the SumLogLR at DT (F_6,78_ = 12.86, p < .00001; see Figure 5.E; BF value above 100). Indeed, Tukey HSD post-hoc tests showed that the SumLogLR at DT was significantly higher at Bin_1_ (1.18 ± 0.06 a.u.) than for any other bin after Bin_4_ (all SumLogLR at DT ≤ 0.84 ± 0.05 a.u.; M1-Disruption and Sham sessions pooled together). Hence, the amount of sensory evidence based on which subjects made their choices decreased as a function of time. Here again, the ANOVA_RM_ did not reveal any significant effect of the factor SESSION (F_6,78_ = 0.04, p = .852) nor of its interaction with the factor BIN (F_6,78_ = 0.41, p = .868). Hence, subjects made their decisions based on a similar amount of sensory evidence in both sessions, suggesting a preservation of the urgency drive during deliberation with M1 disruption.

To further confirm this finding, we obtained a simple approximation of the urgency signal underlying the subjects’ decisions by fitting a linear regression over the rectified version of SumLogLR at DT for each session (M1-Disruption, Sham) and extracted the intercept and the slope of the regression functions (see section *Methods* and Figure 5. F, G and H). Again, student t-tests did not reveal any significant impact of the session on the intercept (t_14_ = 0.48, p =.798) or the slope (t_14_ = −0.22, p =.832) of the urgency functions.

##### Verifying that M1 disruption does not impact deliberation using Bayesian analyses

The ANOVA_RM_ and the t-tests revealed that M1 disruption did not significantly alter the behavioral data measured in the tokens task, suggesting that M1 is not functionally involved in the deliberation process underlying action choices. In order to confirm this result, a Bayes Factor (BF) was computed for each analysis involving the factor SESSION (10 BF values obtained in total), providing us with a ratio of the likelihood probability of the null hypothesis (*i.e.*, H0: the probability that data do not exhibit an effect of SESSION) over the alternative hypothesis (*i.e.*, H1: the probability that data exhibit the effect; Morey and Rouder 2011). A BF value of 1 would reflect an equal probability that H0 and H1 are correct, while a BF value above 1 would reflect a higher probability that H0 is correct. In accordance with conventional interpretation of BF values (Jeffreys, 1961), a BF value ranging between 1 and 3 is interpreted as indicating *anecdotal* evidence in favor of H0, a value between 3 and 10 as indicating *substantial* evidence for H0 and a value between 10 and 30 a *strong* evidence for H0.

Table 1 summarizes the BF values obtained for each factor tested. The average BF value was of 5.66 ± 1.74 (all BFs ranged between 3.46 and 21.28) indicating substantial to strong evidence for an absence of difference in subjects’ behavior between the M1-Disruption and the Sham sessions. Hence, Bayesian analyses further reinforce the conclusion that M1 disruption did not influence the deliberation process underlying decision-making, but solely altered the motor processes that ensue commitment to an action.

**Table 1:** Bayes factor (BF) values. The first column specifies the factors tested for which a BF was computed. Other columns represent the BFs obtained for each behavioral measure in the tokens task. Overall, BFs ranged between 3.46 and 21.28 indicating substantial to strong evidence for a lack of effect of the SESSION on subjects’ behavior.

| Factor tested | DT | %Correct | %TO | SumLogLR | Urgency Intercept | Urgency Slope |
| --- | --- | --- | --- | --- | --- | --- |
| SESSION | 3.87 | 3.77 | 3.91 | 5.23 | 3.46 | 3.48 |
| SESSION*TRIAL | 3.91 | 3.77 | 3.91 | --- | --- | --- |
| SESSION*TIME | --- | --- | --- | 21.28 | --- | --- |

## DISCUSSION

Previous studies have shown that neural activity in motor cortical areas, including M1, is strongly altered during decisions between actions (Alamia et al., 2018; Derosiere et al., 2018; Donner et al., 2009; Gould et al., 2012; Klein-Flügge et al., 2012; Murphy et al., 2016; Steinemann et al., 2018, Thura and Cisek, 2014a, 2016; Tosoni et al., 2008, 2014; Wyart et al., 2012). To date, however, the specific contribution of motor cortical areas to the decision process remains debated. Here, we asked whether M1 causally influences deliberation during action choices, or whether this area mostly contributes to the motor processes overtly expressing commitment. To do so, we tested the impact of a disruption of M1 activity, induced by continuous theta burst stimulation (cTBS), on the behavior of human subjects in *(1)* a simple reaction time (SRT) task allowing us to estimate the duration of the motor processes and *(2)* a modified version of the tokens task (Cisek et al., 2009), which allowed us to estimate subjects’ time of commitment as well as their accuracy criterion.

Subjects were generally faster and more accurate in obvious than in ambiguous trials, suggesting that late decisions relied on weaker sensory evidence compared to early decisions. This is confirmed by the systematic analysis of the sensory evidence available at DT, which indicates that subjects committed to a choice based on less sensory evidence as time elapsed during the course of a trial. This dropping of the accuracy criterion is consistent with previous studies in which similar tasks were used (*e.g.*, Cisek et al., 2009; Gluth et al., 2012; Murphy et al., 2016; Thura et al., 2012, 2014;) and supports recent models postulating that urgency grows over time during speeded decisions (Churchland et al., 2008; Ditterich, 2006; Drugowitsch et al., 2012; Hanks et al., 2011).

The cTBS intervention reduced MEP amplitudes during the entire M1 disruption session, but never following a sham stimulation. Moreover, M1 cTBS lengthened the SRTs (*i.e.*, compared to when sham cTBS was performed), indicating that motor processes that are known to involve M1 were successfully perturbed by the intervention (Huang et al., 2005; Lakhani et al., 2014; McAllister et al., 2013). Notably, in the present study, motor responses were recorded through key-presses. Hence, RTs involved two periods, occurring before and after movement initiation (Spieser et al., 2017). As a consequence, it is sensible to assume that the lengthening of SRT observed here might reflect an increase of *(1)* the time needed for initiating the required motor response, *(2)* the duration of the execution, or *(3)* both.

Critically, all the behavioral data collected in the tokens task were similar in the two cTBS sessions, whether M1 was disrupted or not. Based on this finding, the contribution of M1 to decision-making could be negligible. Hence, past reports of decision-related changes in M1 may reflect the influence of signals coming from upstream structures rather than an actual involvement in the deliberation process itself (Thura and Cisek, 2017; van Maanen et al., 2016). In a similar vein, our recent work shows that M1 disruption negatively alters value-based choices, but only when action values are freshly acquired. Such an effect of M1 disruption does not occur anymore following consolidation. This suggests that M1 contribution to value-based decision-making may vanish as subjects become more proficient at using the value information (Derosiere et al., 2015c; 2017a, 2017b). Thus, in well-learned decision-making tasks, the causal involvement of M1 might be restricted to the motor processes that follow commitment to an action.

However, there are alternative explanations for the lack of effect of M1 disruption on decision behavior in the present study. First, the behavioral variables extracted from the tokens task (*e.g.*, DT, decision accuracy, sensory evidence at DT, *etc.*) may not be sensitive enough to reflect the changes in decision behavior following disruption of motor cortical activity, contrary to the reaction times obtained in the SRT task. In line with this alternative interpretation, the absence of effect of M1 disruption in the tokens task would be due to a lack of sensitivity of the behavioral variables obtained in this specific task. Yet, the results of another study applying microstimulation in the premotor and motor cortex of non-human primates succeeded in altering decision behavior using the same task (Thura and Cisek, 2014b). Hence, this suggests that the behavioral variables extracted from the tokens task are sensitive to the disruption of motor cortical activity.

Also, we cannot rule out the possibility that the cTBS intervention led to some fast reorganization of the decision network following M1 disruption. According to this idea, M1 might still be part of the network involved in the deliberation process but some compensatory mechanisms may have occurred in this network following M1 disruption, leading to similar decision behaviors after M1 and sham cTBS (Bestmann et al., 2004; Briend et al., 2017; Cash et al., 2017; Derosiere et al., 2017a; Rastogi et al., 2017). One way to tackle this issue in the future would be to exploit online rTMS techniques (Duque et al., 2010, 2013), which allow one to perturb neural activity at a specific moment during the decision process, leaving less time for compensatory mechanisms to occur. As such, previous work has shown that while online microstimulation of a decision-related area alters behavior during perceptual decision-making (*i.e.*, the lateral intraparietal area; Hanks et al., 2006), (offline) inactivation of the same area does not (Katz et al., 2016).

Now, if the role of M1 is truly negligible, where in the brain are decisions about actions determined? Among many possible areas, PMd emerges as a promising candidate. First, single-cell recordings in behaving monkeys (Thura and Cisek, 2014a) have shown that during deliberation, activity of some PMd neurons tuned for a particular action reflects the unfolding sensory evidence favoring that action. This observation also makes it possible that this decision-related activity influences M1 neurons through cortico-cortical projections (Duque et al., 2012; Martinez-Gracia et al., 2015). Second, the same studies found that PMd activity related to the selected target reaches a peak about 280 ms before movement initiation regardless of decision difficulty, while a peak of M1 activity occurs about 140 ms later. Third, neurons in the globus pallidus internus, which are insensitive to sensory information during deliberation, become directionally tuned around the time of the PMd activity peak (Thura and Cisek, 2017). Altogether, these results suggest that PMd might be one of the primary sites where decision commitment is determined.

In agreement with this hypothesis, and as mentioned above, a recent study found that microstimulation of PMd neurons alters the deliberation duration, especially if current is applied shortly before commitment time (Thura and Cisek, 2014b). Crucially, stimulation has much less influence on decision duration if it is applied long before commitment or between commitment and movement onset. Relevant for the present work, this study also shows similar time-dependent effect of M1 microstimulation on decision durations - but the effect size is much smaller when M1 is stimulated compared to PMd. Finally, other non-primary motor areas may be causally involved in the deliberation process, including the pre-supplementary motor area (pre-SMA; Tosun et al., 2017). Investigating their precise contribution represents an interesting issue for future investigations.

In conclusion, we show that the offline disruption of M1 activity delays motor processes that follow commitment to an action, but does not alter volitional decision behavior. Taken together, these findings suggest that the contribution of M1 might be downstream of the decision process. Future studies should use online disruption protocols to deal with the putative network reorganization that may have occurred following offline M1 disruption in the present study and broaden their investigation to the role of non-primary motor areas in deliberation, especially the PMd and the pre-SMA.

## GRANTS

This work was supported by grants from the “Fonds Spéciaux de Recherche” (FSR) of the Université Catholique de Louvain, the Belgian National Funds for Scientific Research (FRS-FNRS: MIS F.4512.14) and the “Fondation Médicale Reine Elisabeth” (FMRE). GD was a postdoctoral fellow supported by the FNRS.

## AUTHOR CONTRIBUTIONS

G.D., D.T., P.C. and J.D. conceived and designed research; G.D. performed experiments; G.D. analyzed data; G.D., D.T., P.C. and J.D. interpreted results of experiments; G.D. prepared figures; G.D. and J.D. drafted manuscript; G.D., D.T., P.C. and J.D. edited and revised manuscript; G.D., D.T., P.C. and J.D. approved final version of manuscript.

## DISCLOSURES

No conflicts of interest, financial or otherwise, are declared by the authors.

